# Hsa_circ_0001304 promotes vascular smooth muscle cell autophagy and neointimal hyperplasia through the YTHDF2/mTOR axis

**DOI:** 10.1101/2024.03.13.584799

**Authors:** Shi-Qing Mu, Jia-Jie Lin, Yu Wang, Li-Yun Yang, Sen Wang, Zhao-Yi Wang, An-Qi Zhao, Wen-Jun Luo, Zi-Qi Dong, Yu-Guang Cao, Ze-An Jiang, Si-Fan Wang, Shan-Hu Cao, Li Meng, Yang Li, Shu-Yan Yang, Shao-Guang Sun

## Abstract

**Introduction:** Aberrant autophagy in vascular smooth muscle cells (VSMCs) is associated with the progression of cardiovascular diseases. Recently, circular RNAs (circRNAs) have gradually been reported to regulate autophagy in VSMCs.

**Objectives:** This study aims to investigate the role of hsa_circ_0001304 in VSMC autophagy and its underlying mechanism.

**Methods:** The combined use of dual-luciferase reporter gene assay, MeRIP-qRT-PCR, RIP-qRT-PCR, etc., was performed to verify the regulatory axis, hsa_circ_0001304/miR-636/YTHDF2/mTOR. Cell autophagy detection was performed to uncover the role of hsa_circ_0001304 on VSMC autophagy. The mouse carotid artery ligation model was conducted to assess the role of hsa_circ_0001304 on vascular neointimal hyperplasia.

**Results:** Hsa_circ_0001304 acts as a sponge for miR-636, leading to an increase in the protein levels of YTHDF2. Subsequently, the YTHDF2 promotes the degradation of mTOR mRNA by binding to the latter’s m6A modification sites. Thus, by regulating the miR-636/YTHDF2/mTOR axis, hsa_circ_0001304 activates VSMC autophagy, aggravating neointimal hyperplasia.

**Conclusion:** Our findings suggest that autophagy-related hsa_circ_0001304 could serve as a novel therapeutic target for neointimal hyperplasia-related cardiovascular diseases.

**Including spaces:** Our study documents a novel VSMC autophagy-related circRNA, namely circ-1304, which activates VSMC autophagy through the miR-636/YTHDF2/mTOR axis, thereby exacerbating neointimal hyperplasia. Targeting autophagy-related circ-1304 may contribute to the treatment of neointimal hyperplasia-associated cardiovascular diseases.

## Introduction

Cardiovascular diseases present with a high incidence and mortality rate (Huxley & Perkovic, 2014). Vascular smooth muscle cells (VSMCs) are one of the major cell types that constitute the vascular wall, and their pathological changes are among the primary causes of cardiovascular diseases related to vascular neointima hyperplasia, such as atherosclerosis and hypertension (Newby & Zaltsman, 2000). Physiological cell autophagy is a self-regulating mechanism by which eukaryotic cells degrade damaged proteins or organelles via lysosomes to maintain cellular homeostasis (Devis-Jauregui *et al*, 2021). Importantly, accumulating studies have shown aberrant VSMC autophagy in cardiovascular diseases (Grootaert *et al*, 2018; Tai *et al*, 2016), highlighting the significant role of VSMC autophagy regulation in the treatment of cardiovascular diseases.

Circular RNAs (circRNAs), the new type of regulatory RNA molecules, possess remarkable stability (Kristensen *et al*, 2019). Interestingly, in recent years, two circRNAs, namely hsa_circ_0001402 (Lin *et al*, 2023) and circ-calm4 (Zhang *et al*, 2023), have been identified as regulators of VSMC autophagy, promoting the process. However, more circRNAs involved in VSMC autophagy regulation urgently require investigation.

YTH N6-methyladenosine RNA binding protein F2 (YTHDF2), an m6A binding protein, degrades mRNAs containing m6A sites (Park *et al*, 2019). Notably, YTHDF2 promotes VSMC proliferation and pulmonary arterial hypertension (Hu *et al*, 2023; Qin *et al*, 2021). Mechanistic target of rapamycin kinase (mTOR) is a well-recognized factor known for its ability to inhibit autophagy (Kim & Guan, 2015), including the suppression of VSMC autophagy (You *et al*, 2015). However, it remains unclear whether YTHDF2 regulates the involvement of mTOR in autophagy in an m6A-dependent manner.

Platelet-derived growth factor-BB (PDGF-BB) has been confirmed to induce autophagy in VSMCs (Salabei *et al*, 2013). We previously screened for differentially expressed circRNAs in PDGF-BB-stimulated human aortic smooth muscle cells (HASMCs) (GSE77278), and hsa_circ_0001304 (circ-1304) increased (Chen *et al*, 2020). However, the precise role of circ-1304 in VSMC autophagy remains unclear. In this study, we demonstrate that circ-1304 activates VSMC autophagy, thus aggravating neointimal hyperplasia. The circ-1304/miR-636/YTHDF2/mTOR axis may contribute to the treatment of neointimal hyperplasia.

## Results and Discussion

### Overexpression of circ-1304 promotes VSMC autophagy

Consistent with the microarray data, quantitative real-time PCR (qRT-PCR) analysis revealed an elevation in the expression of circ-1304 in PDGF-BB-stimulated HASMCs (Fig. S1A). Sanger sequencing of qRT-PCR amplified products confirmed that circ-1304 is derived from exons 5-10 of the RBM5 (RNA binding motif protein 5) gene (Fig. 1A), in agreement with the sequence recorded in the circBase database (Glazar *et al*, 2014). Ribonuclease R (RNase R) treatment demonstrated that circ-1304 exhibits strong resistance to degradation, while linear RBM5 mRNA is almost completely degraded (Fig. 1B). Treatment with the transcriptional suppressor actinomycin D indicated that circ-1304 has greater stability and a longer half-life than linear RBM5 mRNA (Fig. 1C). RNase R and actinomycin D analyses further confirmed the circular nature of circ-1304. Nuclear and cytoplasmic RNA detection and RNA fluorescent *in situ* hybridization (FISH) analysis demonstrated that circ-1304 is primarily enriched in the cytoplasm of HASMCs (Fig. 1D-E).

**Fig. 1.**
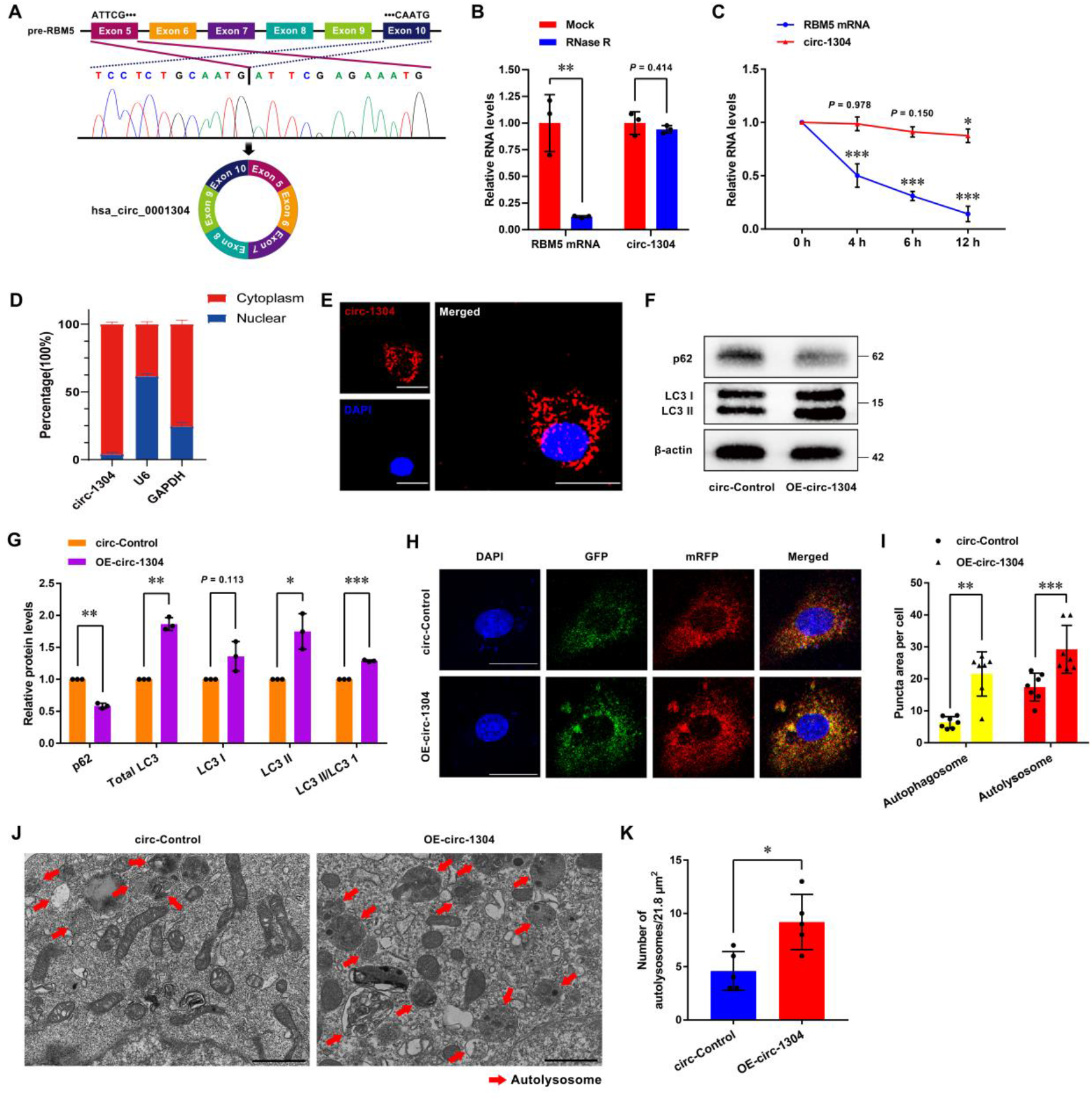
Overexpression of circ-1304 promotes VSMC autophagy. (A) Schematic representation of RBM5 gene exons 5-10 circularization to form circ-1304. (B) qRT-PCR analysis of circ-1304 and RBM5 mRNA levels in total RNA with or without RNase R (3 U/μg) treatment. The RBM5 mRNA served as an RNase R sensitive control. Data are shown as mean ± SD (*n* = 3, ^**^*P* < 0.01 vs. mock group, two-tailed unpaired *t*-test). (C) qRT-PCR analysis of circ-1304 and RBM5 mRNA levels in HASMCs treated with 2 μg/mL actinomycin D. Data are shown as mean ± SD (*n* = 3, ^*^*P* < 0.05, ^***^*P* < 0.001 vs. 0 h group, two-way ANOVA followed by Sidak’s multiple comparisons test). (D) qRT-PCR analysis of the nucleoplasmic distribution of circ-1304 in HASMCs. U6: nuclear positive control; GAPDH: cytoplasmic positive control. Data are shown as mean ± SD (n = 3). (E) RNA FISH analysis of the nucleoplasmic distribution of circ-1304 in HASMCs. Red fluorescence represents the Cy3-labeled antisense probe targeting circ-1304. Blue fluorescence (DAPI) represents the cell nucleus. The scale bar is 12.5 μm. (F-G) Western blot analysis of p62 and LC3 after 48 h of transfection of the expression vector of circ-Control or circ-1304 in HASMCs. Data are shown as mean ± SD (*n* = 3, ^*^*P* < 0.05, ^**^*P* < 0.01, ^***^*P* < 0.001 vs. circ-Control group, two-tailed paired *t*-test). (H-I) Fluorescent images (H) of HASMCs transfected with circ-Control or circ-1304 expression vectors and subsequently infected with mRFP-GFP-LC3 lentivirus for 48 h. The blue fluorescence (DAPI) represents the cell nuclei. The scale bar is 12.5 μm. Quantitative data (I) represents the number of autophagosomes (yellow puncta) and autolysosomes (red puncta) in the merged images. Data are shown as mean ± SD (*n* = 7 HASMCs, ^**^*P* < 0.01, ^***^*P* < 0.001 vs. circ-Control group, Mann Whitney test). (J) TEM analysis of autolysosomes after 48 h of transfection of the expression vector of circ-Control or circ-1304 in HASMCs. The scale bar is 1 μm. (K) Quantify the number of autolysosomes. Data are shown as mean ± SD (*n* = 5, ^*^*P* < 0.05 vs. circ-Control group, two-tailed unpaired *t*-test).

Given the low RNA abundance of circ-1304 in HASMCs, the desired cellular phenotype modulation outcomes could not be achieved through knockdown (Data not shown). Therefore, we employed overexpression (Fig. S1B) to investigate the role of circ-1304 in regulating VSMC autophagy. SQSTM1/p62 (sequestosome 1), a substrate degraded by autophagy (Mizushima *et al*, 2010), was consumed upon overexpression of circ-1304 (Fig. 1F-G). The conversion of MAP1LC3B/LC3 (microtubule-associated protein 1 light chain 3 beta)-I to LC3-II signifies an increase in autophagosome formation (Mizushima *et al*., 2010), and overexpression of circ-1304 promoted this conversion (Fig. 1F-G). The mRFP-GFP-LC3 lentivirus served as an effective tool to monitor autophagic flux (Lin *et al*., 2023). Overexpression of circ-1304 increased the number of autophagosomes (yellow puncta) and autolysosomes (red puncta) in HASMCs (Fig. 1H-I), indicating the activation of autophagic flux. Transmission electron microscopy (TEM) analysis indicated that overexpression of circ-1304 increased the number of autolysosomes in HASMCs (Fig. 1J-K). These results suggest that overexpression of circ-1304 promotes VSMC autophagy.

### Circ-1304 functions through the miR-636/YTHDF2/mTOR axis

Since circ-1304 mainly localizes in the cytoplasm of HASMCs (Fig. 1D-E), coupled with its binding sites with AGO2 protein (Fig. S1C) documented in the circInteractome database (Dudekula *et al*, 2016), we hypothesize that circ-1304 may act as a miRNA sponge. With the help of bioinformatics analysis, we uncovered the intriguing circ-1304/miR-636/YTHDF2 axis. Excitingly, overexpression of circ-1304 led to an elevation in YTHDF2 protein levels (Fig. 2A), while overexpression of miR-636 [achieved through mimic (Fig. S2A)] had the opposite effect (Fig. 2B). Dual-luciferase reporter gene analysis, based on potential binding sites between circ-1304 and miR-636, revealed that the miR-636 mimic attenuated the luciferase activity of the wild-type (WT) circ-1304 plasmid, with no effect on the mutant-type (MUT) plasmid, indicating circ-1304 was a miR-636 sponge (Fig. 2C). RNA FISH analysis further showed that circ-1304 and miR-636 were co-localized in the cytoplasm of HASMCs (Fig. 2D). Dual-luciferase reporter gene analysis, based on potential binding sites between miR-636 and the 3′-untranslated region (UTR) and coding sequence (CDS) region of YTHDF2 mRNA, uncovered that the miR-636 mimic attenuated the luciferase activity of the WT YTHDF2 plasmids, with no effect on the MUT plasmids, indicating the binding of miR-636 to YTHDF2 mRNA (Fig. 2E-F). Circ-1304 indeed enhances YTHDF2 protein levels through competitive binding with miR-636 (Fig. 2G). These findings suggest the presence of the circ-1304/miR-636/YTHDF2 axis.

**Fig. 2.**
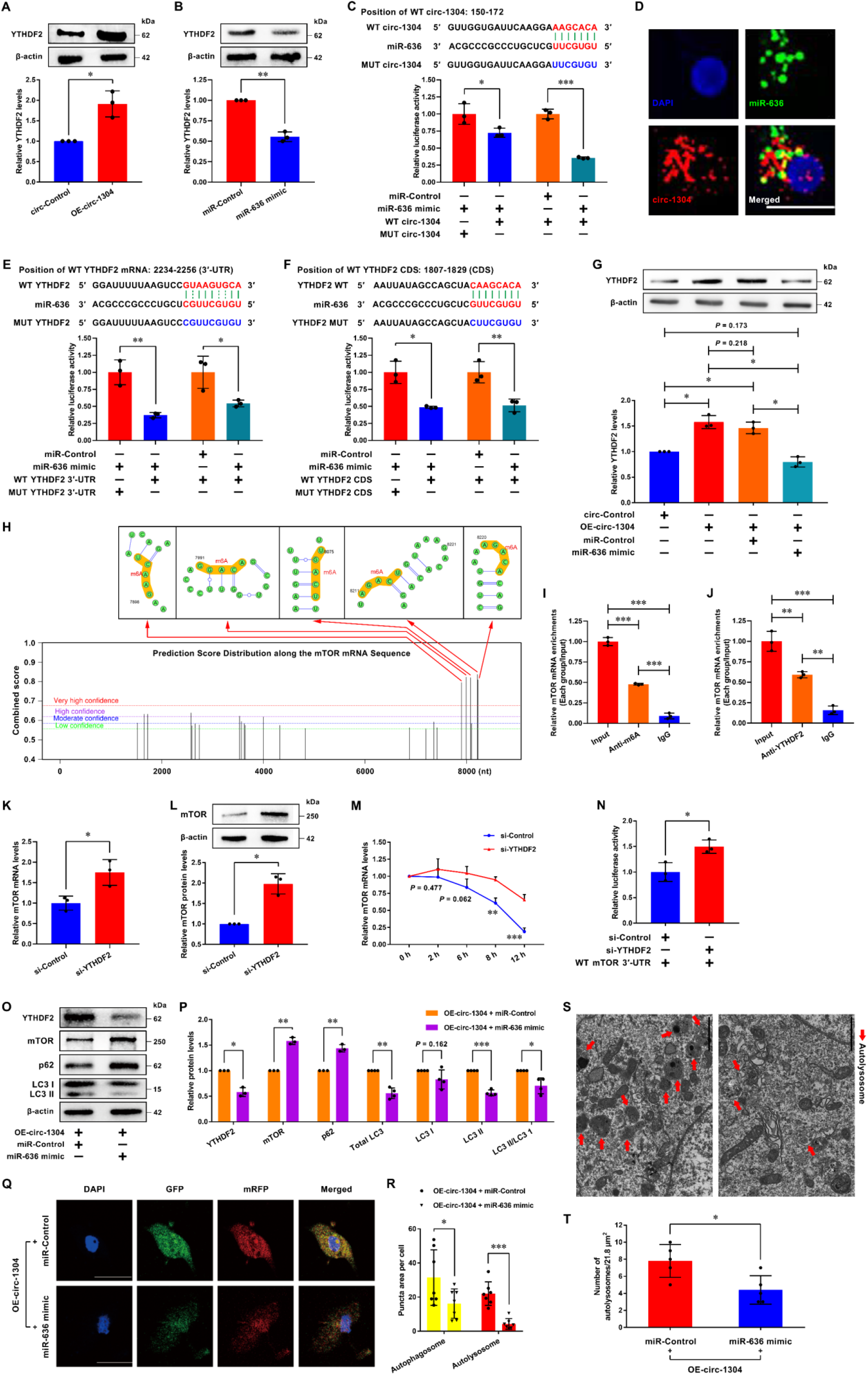
Circ-1304 functions through the miR-636/YTHDF2/mTOR axis. (A) Western blot analysis of YTHDF2 after 48 h of transfection of the expression vector of circ-Control or circ-1304 in HASMCs. Data are shown as mean ± SD (*n* = 3, \**P* < 0.05 vs. circ-Control group, two-tailed paired *t*-test). (B) Western blot analysis of YTHDF2 after 48 h of transfection of miR-Control or miR-636 mimic in HASMCs. Data are shown as mean ± SD (*n* = 3, ^**^*P* < 0.01 vs. miR-Control group, two-tailed paired *t*-test). (C) Dual-luciferase reporter gene analysis of the interaction between circ-1304 and miR-636. WT: wild-type; MUT: mutant. Data are shown as mean ± SD (*n* = 3, ^*^*P* < 0.05, ^***^*P* < 0.001, two-tailed unpaired *t*-test). (D) RNA FISH analysis of circ-1304 and miR-636 co-localization in HASMCs. Red fluorescence represents the Cy3-labeled probe targeting circ-1304. Green fluorescence represents the AF488-labeled miR-636. Blue fluorescence (DAPI) represents the cell nucleus. The scale bar is 12.5 μm. (E) Dual-luciferase reporter gene assay for investigating the interaction between miR-636 and the 3′-UTR of YTHDF2 mRNA. WT: wild-type; MUT: mutant. Data are shown as mean ± SD (*n* = 3, ^*^*P* < 0.05, ^**^*P* < 0.01, two-tailed unpaired *t*-test). (F) Dual-luciferase reporter gene assay for investigating the interaction between miR-636 and the CDS region of YTHDF2 mRNA. WT: wild-type; MUT: mutant. Data are shown as mean ± SD [*n* = 3, ^*^*P* < 0.05, ^**^*P* < 0.01, two-tailed unpaired *t*-test followed by Welch’s correction (left), two-tailed unpaired *t*-test (right)]. (G) Western blot analysis of circ-1304-mediated regulation of YTHDF2 protein levels through competitive binding with miR-636. Data are shown as mean ± SD (*n* = 3, ^*^*P* < 0.05, ne-way ANOVA followed by Tukey’s multiple comparisons test). (H) Uncovering five highly credible m6A modification sites in the 3′-UTR of mTOR mRNA using the SRAMP database. (I) MeRIP-qRT-PCR analysis for the verification of m6A modification sites in mTOR mRNA. Positive and negative controls were established using Input and IgG, respectively. Data are shown as mean ± SD (*n* =3, ^***^*P* < 0.001, one-way ANOVA followed by Tukey’s multiple comparisons test). (J) RIP-qRT-PCR analysis for verifying the interaction between YTHDF2 protein and mTOR mRNA. Positive and negative controls were established using Input and IgG, respectively. Data are shown as mean ± SD (*n* =3, ^**^*P* < 0.01, ^***^*P* < 0.001, one-way ANOVA followed by Tukey’s multiple comparisons test). (K) qRT-PCR analysis of mTOR mRNA levels in HASMCs transfected with si-Control or si-YTHDF2, normalized to GAPDH. Data are shown as mean ± SD (*n* = 3, ^*^*P* < 0.05 vs. si-Control group, two-tailed unpaired *t*-test). (L) Western blot analysis of mTOR after 48 h of transfection of si-Control or si-YTHDF2 in HASMCs. Data are shown as mean ± SD (*n* = 3, ^*^*P* < 0.05 vs. si-Control group, two-tailed paired *t*-test). (M) qRT-PCR analysis of mTOR mRNA levels in HASMCs cells transfected with si-Control or si-YTHDF2 under treatment with 2 μg/mL actinomycin D. Data are shown as mean ± SD (*n* = 3, ^**^*P* < 0.01, ^***^*P* < 0.001 vs. si-Control group at different points in time, two-way ANOVA followed by Sidak’s multiple comparisons test). (N) Dual-luciferase reporter gene analysis of the interaction between YTHDF2 protein and the 3′-UTR of mTOR mRNA containing m6A sites. WT: wild-type. Data are shown as mean ± SD (*n* = 3, ^*^*P* < 0.05, two-tailed unpaired *t*-test). (O-P) Western blot analysis of YTHDF2, mTOR, p62, and LC3 in HASMCs co-transfected with circ-1304 expression vector and either miRNA-Control or miR-636 mimic for 48 h. Data are shown as mean ± SD (*n* = 3 or 4, ^*^*P* < 0.05, ^**^*P* < 0.01, ^***^*P* < 0.001 vs. control group, two-tailed paired *t*-test). (Q) Autophagic flux analysis in HASMCs cells co-transfected with circ-1304 expression vector and either miR-Control or miR-636 mimic for 48 h. The blue fluorescence (DAPI) represents the cell nuclei. The scale bar is 12.5 μm. (R) Quantitative data represents the number of autophagosomes (yellow puncta) and autolysosomes (red puncta) in the merged images. Data are shown as mean ± SD (*n* = 7 HASMCs, ^*^*P* < 0.05, ^***^*P* < 0.001 vs. control group, two-tailed unpaired *t*-test). (S) TEM analysis of autolysosomes in HASMCs co-transfected with circ-1304 expression vector and either miR-Control or miR-636 mimic for 48 h. The scale bar is 1 μm. (T) Quantify the number of autolysosomes. Data are shown as mean ± SD (*n* = 5, ^*^*P* < 0.05 vs. control group, two-tailed unpaired *t*-test).

YTHDF2, an m6A binding protein, recruits the RNase P/MRP complex to degrade mRNAs containing m6A sites. Notably, in an m6A-dependent manner, YTHDF2 facilitates the degradation of PTEN and Hmox1 mRNA, thus promoting VSMC proliferation and exacerbating pulmonary arterial hypertension, respectively. Considering that YTHDF2 degrades target mRNAs containing m6A sites (Park *et al*., 2019), we hypothesized that circ-1304-induced augmentation of YTHDF2 may inhibit well-known autophagy suppressors in an m6A-dependent manner, thereby promoting VSMC autophagy. Through bioinformatics analysis, the autophagy inhibitor mTOR (Kim & Guan, 2015) has come into our focus. Utilizing the SRAMP database (Zhou *et al*, 2016), we observed the presence of five high-confidence m6A modification sites on the 3′-UTR of mTOR mRNA (Fig. 2H). MeRIP-qRT-PCR analysis validated the existence of m6A modification on mTOR mRNA (Fig. 2I), and RIP-qRT-PCR analysis demonstrated the ability of YTHDF2 protein to bind to mTOR mRNA (Fig. 2J), indicating that YTHDF2 protein interacts with mTOR mRNA in an m6A-dependent manner. Furthermore, the knockdown of YTHDF2 [achieved through siRNA (Fig. S2B)] resulted in elevated expression levels of both mTOR protein and mRNA (Fig. 2K-L). Analysis using actinomycin D revealed that the knockdown of YTHDF2 impeded the degradation of mTOR mRNA (Fig. 2M). Dual-luciferase reporter gene assay indicated that the knockdown of YTHDF2 increased the luciferase intensity of the WT mTOR mRNA plasmid (Fig. 2N). These findings suggest that YTHDF2 protein inhibits mTOR expression in an m6A-dependent manner. mTOR is well-established factor inhibiting autophagy. Interestingly, knockdown of mTOR mRNA activates VSMC autophagy.

Subsequent rescue experiments were conducted. The introduction of miR-636 rescues VSMC autophagy induced by circ-1304 overexpression by decreasing YTHDF2 levels and elevating mTOR levels, thus inhibiting VSMC autophagy (Fig. 2O-T).

Collectively, these findings indicate that circ-1304 promotes VSMC autophagy through the miR-636/YTHDF2/mTOR axis.

### Overexpression of circ-1304 aggravates neointimal hyperplasia

In a mouse model of neointimal hyperplasia induced by common carotid artery ligation, the *in vivo* role of circ-1304 was investigated. Following *in situ* delivery for 14 days, lentivirus-mediated overexpression of circ-1304 resulted in exacerbated neointimal formation (Fig. 3A-B). Furthermore, changes in the expression levels of autophagy-related proteins were observed, including increased YTHDF2 and total LC3 levels (Fig. 3C, D, I, and J), and decreased mTOR and p62 levels (Fig. 3E-H). These findings suggest that overexpression of circ-1304 exacerbates neointimal hyperplasia by enhancing YTHDF2 protein levels and suppressing mTOR protein levels.

**Fig. 3.**
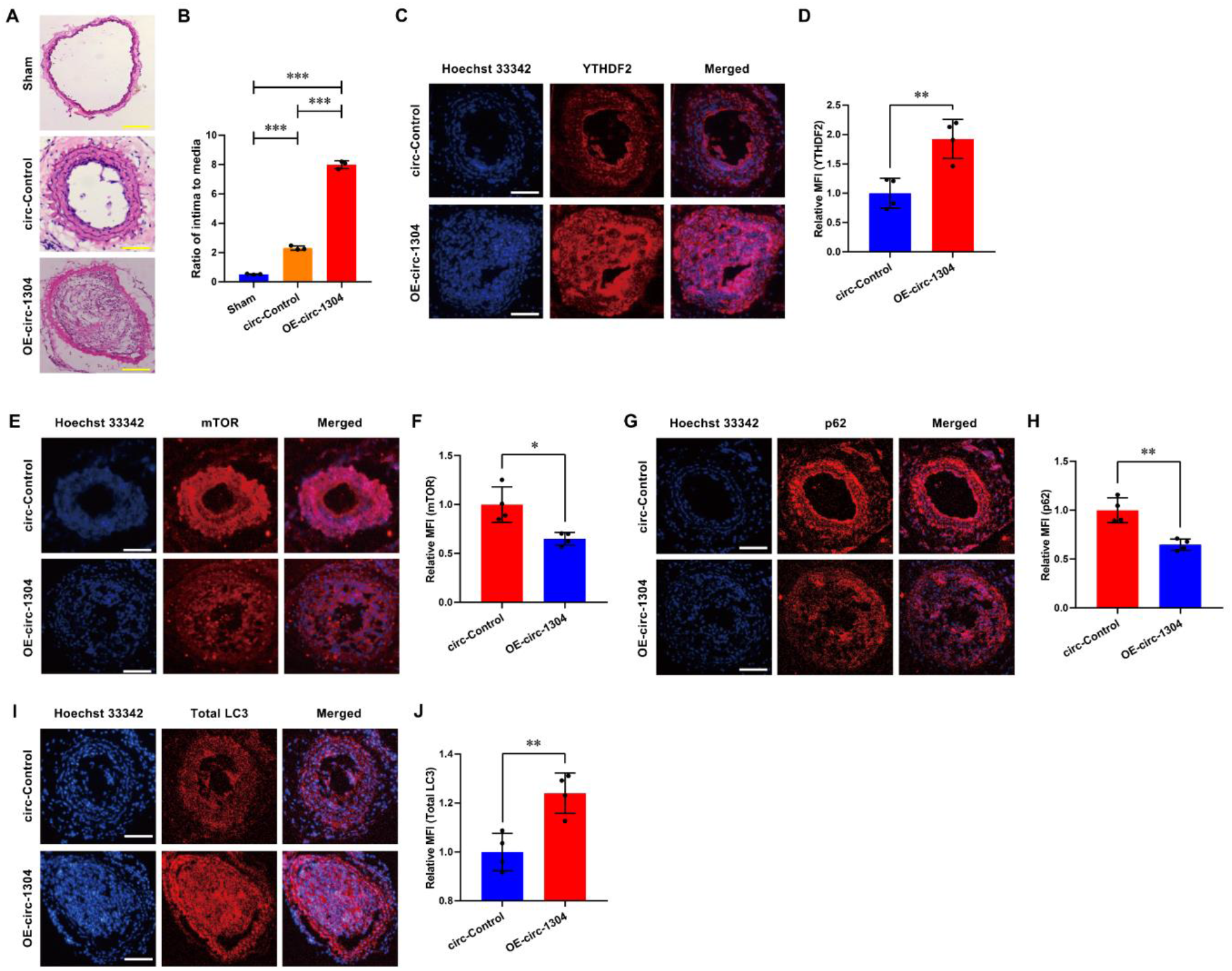
Overexpression of circ-1304 aggravates neointimal hyperplasia. (A) The cross-sectional hematoxylin and eosin (H&E) staining images of the ligated mouse common carotid arteries after 14 days of treatment with Sham Control, circ-Control lentivirus, or circ-1304 lentivirus. The scale bar is 100 μm. (B) Morphometric measurement of the intimal/media ratio in the mouse common carotid arterial sections (*n* = 3 mice). Data are shown as mean ± SD (^***^*P* < 0.001, one-way ANOVA followed by Tukey’s multiple comparisons test). (C-J) Immunofluorescence of YTHDF2, mTOR, p62, and total LC3 after 14 days of *in situ* delivery of circ-Control or circ-1304 lentivirus in ligated mouse common carotid arteries. The scale bar is 100 μm. The quantification of YTHDF2, mTOR, p62, and total LC3 levels in the ligated mouse common carotid arteries (*n* = 4 mice). Data are shown as mean ± SD (^*^*P* < 0.05, ^**^*P* < 0.01 vs. circ-Control lentivirus group, two-tailed unpaired *t*-test). MFI: mean fluorescent intensity.

## Methods

Detailed material and methods are available in the supplementary data.

### Ethics statement

All animal experiments carried out adhered to the ARRIVE guidelines and received approval from the Institutional Animal Care and Use Committee of Hebei Medical University (Approval no. 20180104).

### Ligation model and lentivirus infection of mouse common carotid arteries

Male C57BL/6J mice aged eight weeks were obtained from the Experimental Animal Center at Hebei Medical University (Shijiazhuang, China). Neointimal hyperplasia was induced by ligating the common carotid artery in mice, as described in previous studies (Kumar & Lindner, 1997; Lin *et al*., 2023). The laboratory where the mice were housed maintained a pathogen-free environment with a 12-hour light/dark cycle at a temperature of 22℃, and provided unlimited access to food and water. Pluronic gel was effectively used as a medium for *in situ* RNA delivery (Lin *et al*., 2023; Smolock *et al*, 2012; Sun *et al*, 2011). Regarding circRNA delivery, 20 μL of circ-Control or circ-1304 lentivirus (1 × 10^9 TU/mL), 5 μg/mL Polybrene (Sigma-Aldrich), and 8 μL of GP-transfect-Mate transfection reagent (GenePharma) were incubated to prepare a 30 μL of 20% F-127 pluronic gel (Sigma-Aldrich) at 4℃ for 2 h. Anesthesia using 2% isoflurane was administered to the mice before performing ligation near the carotid bifurcation on the left common carotid artery. Subsequently, 30 μL of pluronic gel containing the respective RNA for in situ delivery was immediately applied to each exposed artery. The sham group underwent a procedure in which sutures were placed around the exposed arteries but untightened. The surgery duration for each mouse was approximately 20 min. After two weeks following the surgical procedure and RNA treatment, as outlined previously (Lin *et al*., 2023; Zhu *et al*, 2019), the mice were sacrificed using an excessive amount of isoflurane. The ice-cold PBS was perfused through the left ventricles to obtain vascular tissues for hematoxylin and eosin (H&E) staining, as well as immunofluorescence staining. A single operator conducted all the procedures, and each group was comprised of six mice randomly assigned.

### Statistical analysis

The data presented in this study represents the mean ± standard deviation (SD) of at least three independent experiments. Statistical analysis was performed using the paired *t*-test for paired samples with differences following a normal distribution. For unpaired samples, the Mann-Whitney test was utilized when the data did not conform to normality, the unpaired *t*-test was chosen if the data met the assumptions of normality and homoscedasticity, the unpaired *t*-test with Welch’s correction was applied in situations where the assumption of normality was met but not homoscedasticity. For multiple groups of samples with a single factor, the one-way ANOVA followed by Tukey’s multiple comparisons test was performed if the data met the assumptions of normality and homoscedasticity. For multiple groups of samples with two factors, the two-way ANOVA followed by Sidak’s multiple comparisons test was performed if the data was normally distributed. A significance threshold of *P* < 0.05 was set.

## Acknowledgments

This work was supported by the National Natural Science Foundation of China (grant nos. 82170439, 81670273, and 81200215 to S.-G.S.), the Natural Science Foundation of Hebei Province (grant no. H2021206399 to S.-G.S.), and the Natural Science Foundation of Beijing (grant no. 7242013 to S.-Y.Y.).

## Author Contributions

**Shao-Guang Sun:** conceptualization, resources, supervision, funding acquisition, methodology, project administration, writing – review & editing. **Shu-Yan Yang:** conceptualization, supervision, funding acquisition, methodology, project administration, writing – review & editing. **Shi-Qing Mu:** data curation, software, investigation, formal analysis, validation, visualization, writing – original draft. **Jia-Jie Lin:** data curation, software, investigation, formal analysis, validation, visualization, writing – original draft. **Yu Wang:** data curation, software, investigation, validation, visualization, writing – original draft. **Li-Yun Yang:** investigation, validation. **Ze-An Jiang:** investigation, validation. **Shan-Hu Cao:** investigation, visualization. **Zhao-Yi Wang:** investigation, validation. **An-Qi Zhao:** investigation, validation. **Sen Wang:** investigation, validation. **Si-Fan Wang:** investigation. **Wen-Jun Luo:** investigation. **Zi-Qi Dong:** investigation. **Yu-Guang Cao:** investigation. **Li Meng:** investigation. **Yang Li:** investigation.

## Disclosure and competing interests statement

The authors declare no competing interests.

## Data Availability Section

This study includes no data deposited in external repositories.

